# At the roots of interference control: Conflict task in rats reveals the commonalities of onto- and phylo-genetic development

**DOI:** 10.1101/2024.05.22.595426

**Authors:** Julien Poitreau, Frédéric Ambroggi, Thierry Hasbroucq, Francesca Sargolini, Boris Burle

## Abstract

Responding to internal goals and filtering irrelevant environmental cues is essential for adapted behavior. To better understand their phylogenetic evolution, we explored the underlying mechanisms in rats, thanks to the adaptation of a well-established conflict task in Humans. Besides mean performance, state of the art data analysis based on distribution analysis and formal modeling (diffusion model adapted to conflict tasks that proved very powerful in Humans) revealed mechanisms similar to the ones observed in human adults, grounding any theoretical explanation into an evolutionary perspective. Besides, the dynamics of the underlying processes in this simple conflict task resembles the ones observed in human children performing a more challenging conflict task, including the tendency to loose goals. The present study bridges the gap between onto- and phylo-genetic development of cognitive control, opening new perspectives to understand both their functional and neural implementation.

## Introduction

Adapted behaviors require to appropriately select relevant actions and to refrain from responding to irrelevant environmental cues. How inhibitory control developed through the phylogeny is a core question to understand the evolution of cognitive abilities (MacLean et al., 2014). The capacity to inhibit irrelevant actions and to perform the relevant ones is classically studied in so-called “conflict tasks”, such as the Stroop task (Stroop, 1935), the Simon task (Simon & Small, 1969) and the Eriksen flanker task (Eriksen & Eriksen, 1974). For example, in a typical version of the Simon task, participants must issue a left or right response as a function of the color of the stimulus, which can be presented on the left or on the right side of a reference point. Although entirely irrelevant for the task at hand, stimulus position interferes with color processing and must be suppressed to provide the correct response, leading to degraded performance (slower response time – RT – and higher error rate, often referred to as “Simon effect”) when the position of the stimulus and the response do not match (incompatible trials) compared to when they do (compatible trial). The simplicity of the experimental design (see Hommel, 2011, for an overview) makes the Simon task especially fitted to investigate response selection processes beyond young adults. It is, indeed, easily transferable to young children (see *e*.*g*. Ambrosi, Servant, Blaye, & Burle, 2019; Ambrosi, Śmigasiewicz, Burle, & Blaye, 2020; Iani, Stella, & Rubichi, 2014; Smigasiewicz, Ambrosi, Blaye, & Burle, 2020), but also to non-human animals: a Simon effect has been reported in non-human primates (Huguet, Barbet, Belletier, Monteil, & Fagot, 2014; Nakamura, Roesch, & Olson, 2005), in rats (Courtière, Hardouin, Burle, Vidal, & Hasbroucq, 2007; Courtière et al., 2011), in pigeons (Urcuioli, Vu, & Proctor, 2005), and even in ants (Czaczkes, Berger, Koch, & Dreisbach, 2022). These demonstrations of a Simon effect at different levels of the phylogeny might impose constraints on the theoretical explanation of the underlying processes and on the evolution of cognitive control: Indeed, if the Simon effect is driven by the same processes on all those species, any explanation of the effect (including in humans) must be compatible with the currently known cognitive abilities of all species presenting the effect. However, the hypothesis that the same processes underlie the observed effects in all species is not mandated, as convergent evolution may lead to similar behavioral effects but driven by very different underlying processes, both across species (*e*.*g*. Dépy, Fagot, & Vauclair, 1997), and during development in humans (see Chi, 1977, for a discussion).

A key characteristic of the Simon effect in humans is its dynamics: it is a transient, short-lived effect (van den Wildenberg et al., 2010, for an overview), as revealed through distribution analysis of errors and RTs. Besides mean error rate, the so-called Conditional Accuracy Functions (CAF, Gratton, Coles, Sirevaag, Eriksen, & Donchin, 1988; Lappin & Disch, 1972, see methods), reveals that for incompatible settings, the probability of a correct response drops dramatically for fast responses, and reaches a plateau around almost 100% correct for slow responses (*e*.*g*. van den Wildenberg et al., 2010). In contrast, it is almost flat, close to ceiling, on compatible trials. In terms of RT distribution in humans, the difference between incompatible and compatible trials is large for fast responses, but decreases, and tends to disappear, for slow responses (Pratte, Rouder, Morey, & Feng, 2010; Ridderinkhof, 2002). Such distributional approaches have seldomly been used in non-human animals. Huguet et al. (2014) analyzed both RT distributions and CAFs in Baboons (*papio papio*), and revealed dynamics comparable to the one observed in humans. In rats, Courtière et al. (2007) reported a slight reduction of the Simon effect (with auditory stimuli) for long RTs, but did not present CAF analysis, leaving open the question as to whether the Simon effect really has the same dynamics as in humans. The main goal of the present study was to assess whether the processes at hand in rats performing a Simon task are similar to the ones evidenced in humans by analysing the underlying dynamics.

To reach this goal, rats were trained in an adapted Simon task (Figure 1, see Methods for detailed procedure). Performance was analyzed with distribution analyses and a process model of conflict tasks developed for humans (Ulrich, Schröter, Leuthold, & Birngruber, 2015) was fitted to those data to even better assess the dynamics of the underlying processes.

**Figure 1.**
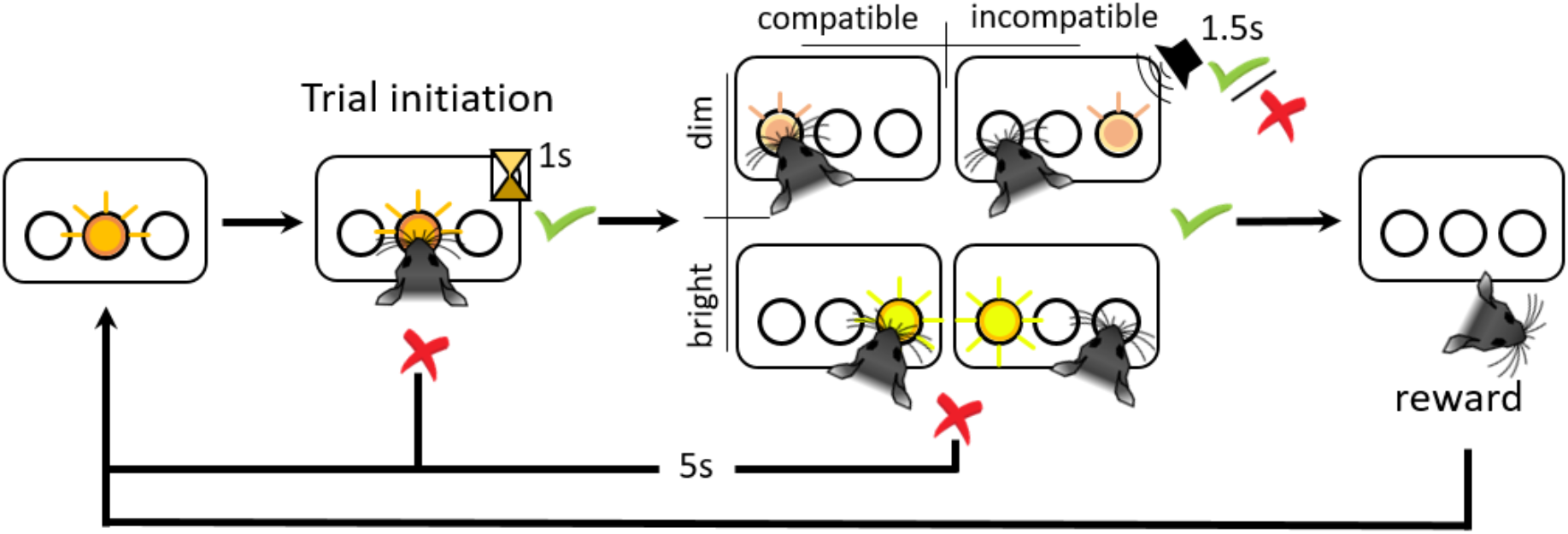
Adapted Simon task in rats. Animals have to respond with a right or left nose poke to the intensity of a light stimulus (*e*.*g*. Bright stimulus – right response; Dim stimulus – left response). The response rule (intensity – response side) is counterbalanced across animals. Stimulus position is irrelevant for the task and has to be ignored. Trial initiation implies a 1s maintained central poke. Stimulus (dim or bright) appears for 500ms. Animals have then 1.5s to respond in one of the two lateral ports as a function of intensity. End of the response window is indicated by an auditive feedback differing as a function of response correctness. An incorrect response lead to a 5s punishment delay with all the lights off. A poke in the feeder after a correct response triggers the reward (sugar pellet).

## Methods

All data and processing scripts can be found at

https://osf.io/pz7m8/?view_only=1250bfdc3c654bc3aaf75b23ca87da9f.

### Animals

23 adult male Long Evans rats (250g at the beginning of the behavioral procedure) were used for this experiment. Animals were housed in pairs in standard cages and kept in a temperature- (21*°*C) and hygrometry-controlled room, under a 12h dark-light cycle. All experimental procedures were done during the light phase. Animals were kept at 85% of their normal body weight throughout the entire behavioral protocol. They had unlimited access to water in their home cages. All experimental procedures were approved by the local ethical committee and the French authority under the reference number: APAFIS#9751-2017042712424200.

### The Simon Task

The adapted Simon task is depicted on Fig. 1 (see supplementary methods for details on the shaping procedure). Animals learned to associate a light stimulus of two different intensities, dim or bright, to a left or right side response (poke into one of two lateral nose ports, NP). Twelve animals learned the association “bright stimulus – left-side response” and “dim stimulus – right-side response” while 11 learned the opposite one. Although irrelevant for the task at hand, the stimuli could be presented either on the left or on the right of the central NP. This led to two types of trials derived from stimulus position and intensity: compatible trials when rats had to respond on the same side as stimulus position; incompatible trials when they had to respond on the opposite side. The temporal organisation of trial events was as follows: Central NP was first illuminated indicating the possibility to initiate a trial (central light intensity was of middle intensity between the dim and the bright stimuli). Animals had then to hold their nose into the central NP for at least 1s. After this holding delay, the light stimulus (bright or dim) appeared for 0.5s inside either the left or right NP. From then on, animals had 1.5 s to complete their response after stimulus onset. They had to poke with their nose either on the left or the right NP as a function of stimulus intensity and regardless of the presentation side of the stimulus. At the end of the response window, an auditory feedback (low or high pitch) indicated response correctness. Following a correct response, rats had then 5s to poke in the feeder to obtain a sucrose pellet (45mg, dustless; TestDiet®, St Louis, Missouri).

After pellet collection, the next trial could be initiated. If the 1s central NP holding time was not respected, the trial did not start, and the animal had to start a new trial. No response before the end of the response window was scored as an omission. In case of wrong response or omission, the same type of trial was re-started after a 5s punishment delay, during which all the NPs were off. Repeating erroneous trials was used to avoid stimulus independent stereotypical responses on the same lateral NP. In case of several correct responses in a row, stimulus position and intensity were pseudo-randomly distributed to prevent more than three repetitions of the same type of trial. Sessions were 30min long or ended after animals collected a maximum of 150 pellets. Analysed data were acquired during 16 sessions (8 consecutive days) after a performance plateau was reached.

## Data analyses

### Pre-processing

All task events (i.e. stimuli, pokes, rewards), along with their absolute times, were recorded during 16 test sessions (temporal precision: 10 ms). Data were first extracted with MedPC-to-Excel utility and then processed with Python and R scripts that are accessible at xxxx). For each compatibility condition, response time was measured as the time between the stimulus apparition and the poke in the lateral NP, and accuracy was calculated as the number of correct responses over the total number of responses. Median absolute deviation (MAD) of the response times distribution (Leys, Ley, Klein, Bernard, & Licata, 2013) was calculated for each rats in each compatibility condition. Data superior or inferior to 2.5 times the MAD was considered outliers and excluded from the analysis (about 0.12% of the data). Anticipations, defined as response times under 300 ms (about 11%), and omissions (about 14%) were also excluded from analysis.

### Mean performance analysis

We first analysed mean performance with canonical repeated measures analyses of variances with compatibility as within-participant factor. For accuracy, these analyis were performed on arcsine square root transformed data to stabilize the variances.

### Distribution analysis

Besides mean effects, we analyzed the dynamic of performances as a function of response times, through response times distribution analysis. Response times of compatible and incompatible trials for each rat were vincentized (Ratcliff, 1979; Vincent, 1912) for all 16 sessions: briefly, RT were sorted in ascending order and binned into 7 bins of equal size, and the means for each bin were computed. These mean values per bin were then averaged across rats to obtain a “mean” cumulative density function, which is representative of the individual ones (Rouder & Speckman, 2004). The magnitude of the Simon effect as a function of response times was then measured as the difference between the incompatible and compatible bins. Plotting these differences as a function of response time, is often referred to as the “delta-plots” (De Jong, Liang, & Lauber, 1994; Pratte et al., 2010; Ridderinkhof, 2002). Distribution analysis were also performed on the proportion of correct responses for each type of trials. As previously, RTs were vincentized and the proportion of correct responses within each bin was computed. Plotted against response times, these proportions allow to obtain the conditional accuracy function (CAF).

The dynamic of the Simon effect on response times and accuracy as a function of response times was also assessed with repeated measures ANOVAs with compatibility and quantiles as within-participant factors. To better characterise the shape of the CAF, polynomial contrats analysis were performed (see Ambrosi et al., 2020; Burle, Spieser, Servant, & Hasbroucq, 2014): both linear and quadratic components were analysed.

### Gaussian Mixture analysis

The CAF obtained led us to explore more precisely the shape of the correct and incorrect RT distributions, from which we suspected the presence of two different mixed distributions contributing to the observed RT ones.

We therefore performed Gaussian mixture analysis with scikit-learn (Pedregosa et al., 2011) function `GaussianMixture` from the `mixture` module. RT distribution being notably right-skewed, they were first log-transformed to be closer to Gaussian distributions. In order to reduce the number of parameters to be fitted, we assumed an equal variance for the two mixed distributions. Single distribution models *vs*. two distributions ones were compared based on the Akaike Information Criterion (AIC, Akaike, 1974). The selected model was back-transformed into real time (in ms) by an exponential transformation to allow direct comparison with empirically obtained distributions. Such mixture analysis was performed separately for correct and error RT distributions of both compatible and incompatible conditions for all rats. The large majority of analyses (80 over 92) favored the mixture model based on the AIC criterion. For the cases where a single distribution was favored, the two models were actually very close (see Fig. S1). To keep these few cases in the analysis, we hence decided to force the mixture analysis to find two distributions, which led to visually satisfactory solutions on all cases (see supplemental online material). The model provided parameters for both distributions (log-transformed). To better analyse the first distribution (D1, which should correspond to the “real RT”, see results), we used the parameters to simulate data corresponding to those parameters, in order to proceed to the same analyses as experimental data and compared the changes in the results given the considered distributions. We randomly generated sampled RTs stemming from each distribution, in a proportion corresponding to the actual proportion of the corresponding trials in the condition and rat under analysis (overall 10,000 trials were generated per rat and compatibility conditions). The obtained data were back-transformed into real time (in ms) by an exponential transformation.

### DMC modeling

Capitalizing on the success of “Accumulation to bounds” models, Ulrich et al. (2015) proposed a generic model of conflict tasks, the “Diffusion Model for Conflict” (hereafter named “DMC”), that proved successful in accounting for human adult data in the Eriksen-Flanker task (Ulrich et al., 2015), and the Simon task (Servant, White, Montagnini, & Burle, 2016; Ulrich et al., 2015), and efficiently models speed-accuracy trade-off in such tasks (Mittelstädt, Miller, Leuthold, Mackenzie, & Ulrich, 2022). The model also proved to nicely fit children data, providing a better understanding of developmental trends in the Eriksen-flanker, the Simon and the Stroop task (Ambrosi et al., 2019). To the best of our knowledge, this model has never been fitted to non-human data. Would the model correctly account for the data obtained in rats, this could have important consequences. First, being a process model (*i*.*e*. that aims to model the processes leading to the observed data), a good fit would strengthen the idea that the same processes are at stake in both humans (adult and children) and rats. Besides, it could provide more insights on the dynamic of such underlying processes allowing comparison with human. The model was fitted with python version of the pyDMC package (Mackenzie & Dudschig, 2021)^1^. It is not the place to describe in details the model (see Ulrich et al., 2015); only essential aspects will be presented here. Briefly, the model assumes two processes: a transient automatic activation of the response, whose strength is capture by the “*A*” parameter and that reaches it peaks at “*Lat*” latency (see below for more details) and a controlled process described as a standard diffusion model with key components being the rate of information accumulation (the drift rate, “*μ*”) and the threshold for the response to be emitted (“*b*”). The two processes feed a single evidence accumulator. Besides the decision processes, the whole RT is obtained by adding a residual time whose mean is “*Ter*”.

## Results

### Mean performance

Percentage of correct responses was lower for incompatible trials (73.3 %) than for compatible ones (91.9 %, [F(1,22) = 82.87; *p <* .0001]). RT were longer for incorrect trials (578 ms) than for correct ones (511 ms), and this factor interacted with compatibility [F(1, 22) = 5.71 *p <* .026]. A clear compatibility effect was present for correct responses (489 vs 532 ms, for compatible and incompatible respectively, [F(1, 22) = 29.57 *p <* .0001]), but not for errors [F(1, 22) *<* 1]).

## Distribution analyses

### Accuracy

Repeated measures ANOVA on CAF (Figure 2 panel A) revealed a main effect of compatibility [F(1, 22) = 82.9; *p <* .0001], of quantiles [F(6, 132) = 23.0; *p <* .0001], and an interaction between the two [F(6, 132) = 2.88; *p <* .01]. Trend analysis (orthogonal polynomial, Ambrosi et al., 2020; Burle et al., 2014) revealed both a linear (negative going trends) and a quadratic (convex trends) component [*β*= −0.07, F(6, 132) = 34.90; *p <* .0001 and *β*= −0.09, F(6, 132) = 52.66; *p <* .0001 for linear and quadratic respectively] on compatible trials. For incompatible trials, no linear trend was revealed [*β* = −0.01, F(6, 132) = .56], while a quadratic one was observed [*β*= −0.12, F(6, 132) = 78.31; *p <* .0001], indicating an almost perfect inverted U-shape.

**Figure 2.**
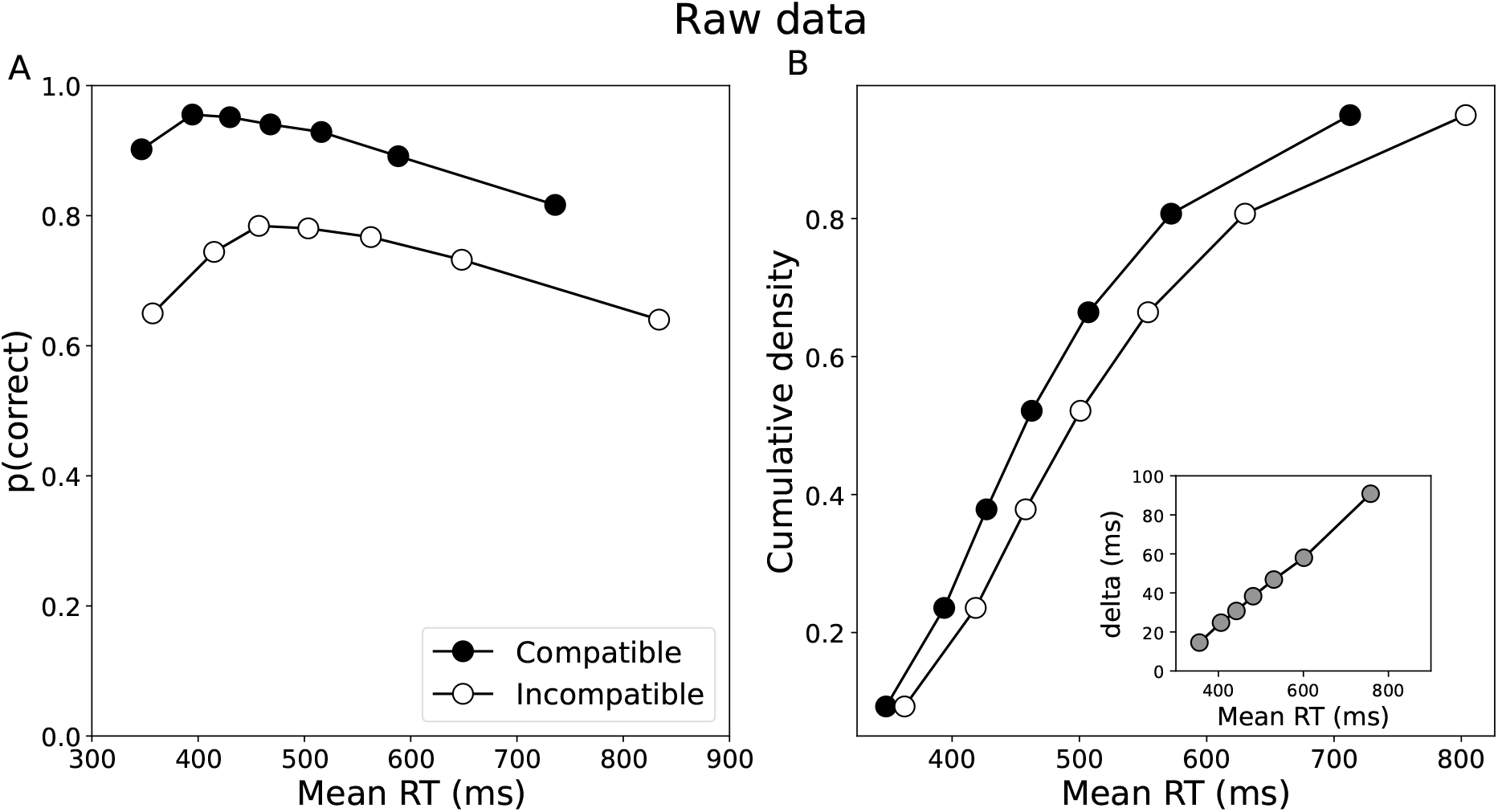
Distribution analysis. A. Conditional Accuracy Function (CAF) for compatible (black circles) and incompatible (open circles) trials. A decrease in accuracy for fast RT for incompatible trials, is followed by asubsequent decreases for long RT. B. Cumulative Density Functions (CDF) of the correct responses. Inset: mean interference effect (incompatible - compatible) as function of RT. The size of the interference effect increases linearly with RT.

### Response times

Repeated measures ANOVA on the cumulative density function of response times (Figure 2 panel B) revealed a main effect of compatibility [F(1, 22) = 29.57; *p <* .0001], a trivial main effect of quantiles [F(6, 132) = 241.58; *p <* .0001], and an interaction [F(6, 132) = 10.44; *p <* .0001], which, supported by the slope of the delta plots, indicates an increase of the interference effect as response times became longer.

### Mixture analysis

To clarify the origin of the performance decrease for long RT revealed by the CAF, we analysed in more detail the shape of the RT distributions through a kernel density estimates (Figure 3). For short RTs, there were more errors than correct responses accounting for the initial drop of performance. Besides these fast errors, a clear bimodality was evident in the incorrect trials distribution, showing an increase in the number of errors for slow responses. To better assess this bimodality, mixture analyses were performed (see Methods for more details), for each rat and all four distributions (correct and incorrect responses, for compatible and incompatible trials). For all rats, and in all four cases, a first recovered distribution (hereafter termed D1) contained rather “fast” RT, and was right skewed. The second distribution (hereafter termed D2) was much slower with a dynamics that departs from standard RT distributions (see detailed parameters of both distributions in Table 1 in supplementary online material). Although present in both correct and error trials, we evaluated whether this “non-RT” distribution could potentially account for the decrease in accuracy for long RTs observed in the CAFs. If such is the case, removing these slow trials from both correct and errors RT distributions should remove the decreased accuracy for long RTs. To test this proposition, we randomly generated sampled RTs stemming from each distribution (see SM for more details) and processed these generated “RTs” in the same way as experimental data. Only D1 results are presented here (see section in Supplementary online material for D2 results). Repeated measures ANOVA revealed a significant compatibility effect on mean accuracy [F(1,22) = 162.54; *p <* .0001] and on mean RT (compatible: 419 ms, incompatible: 451 ms, (1,22) = 37.50; *p <* .001]. They were alse shorter for errors than for correct trials (correct: 444 ms, errors: 425 ms, F(1,22) = 14.12; *p <* 0.002), with no interaction effect between these two factors [F(1,22) = 0.08; *p* = .78].

**Table 1.** Parameters of the DMC for the current study, along with estimates from human studies, for sake of comparison. Meaning of parameters: A is the amplitude (i.e., the strength) of the automatic activation, and Lat is the time at which it reaches its peak. μ corresponds to the mean accumulation information rate in the rule-based processing route. The threshold at which the decision is made correspond to the b parameter. The T_er_ captures the non decision components of the RT (stimulus encoding and response execution).

| Study | Population | parameters |  |  |  |  |
| --- | --- | --- | --- | --- | --- | --- |
| | | A | Lat | $\mu$ | b | Ter |
| Current | rats | 18.4 | 134 | 0.44 | 38.3 | 376 |
| Ambrosi et al., 2019 | Children | 28.8 | 54 | 0.36 | 80.6 | 532 |
| Ulrich et al., 2015 | Adults | 16.0 | 35 | 0.69 | 54.6 | 323 |
| Servant et al. 2016 | Adults | 15.1 | 29 | 0.47 | 60.5 | 335 |
| Mittelstädt et al. 2022 | Adults |  |  |  |  |  |
| Speed |  | 25.6 | 40 | 0.48 | 47.7 | 305 |
| Accurate |  | 21.8 | 57 | 0.70 | 72.3 | 343 |

**Figure 3.**
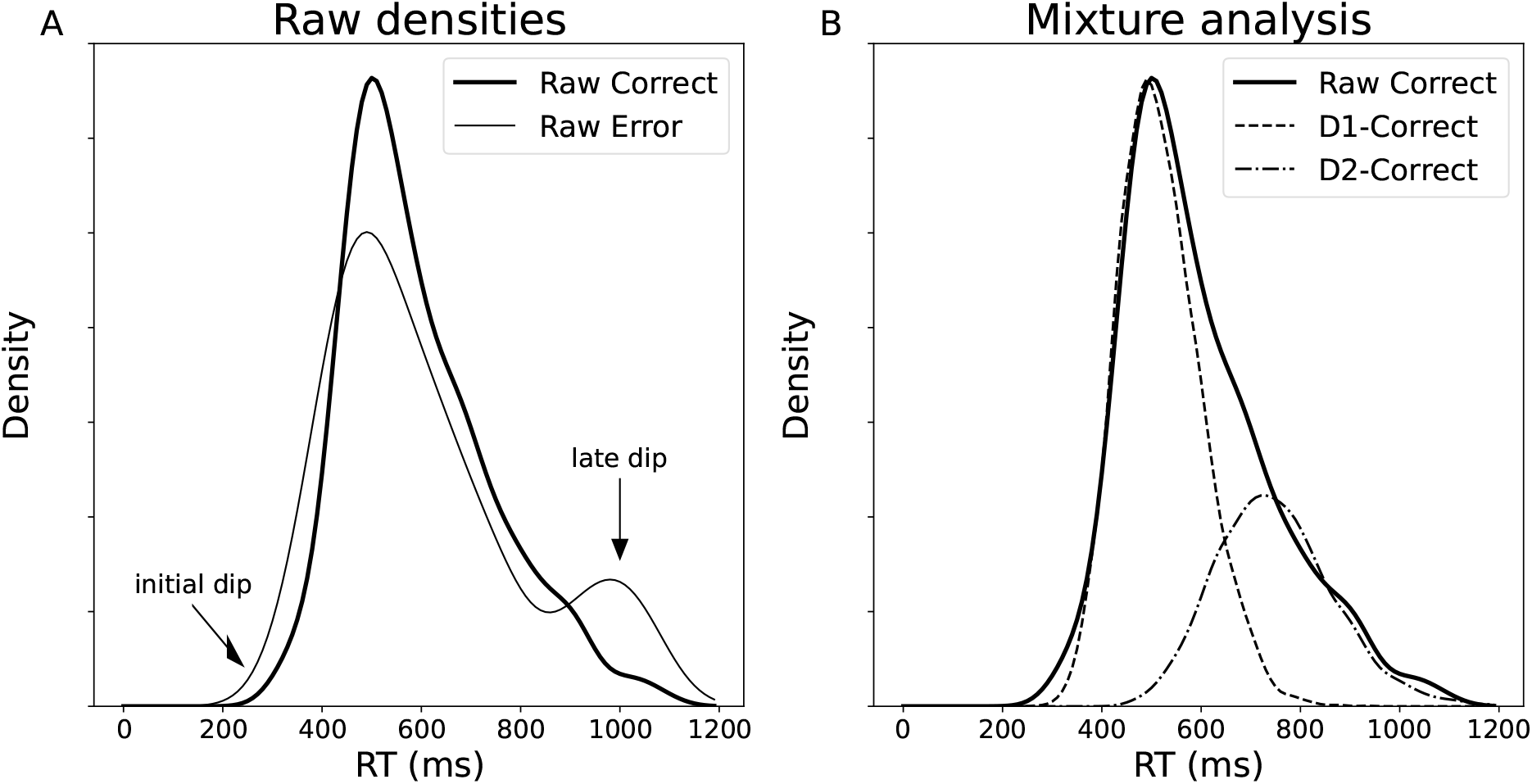
Distribution shape analysis through Kernel Density Estimate (kde) and mixture analysis. Panel A: Example of kde for “Correct” (thick line) and “Error” (thin line) for a given rat in a given condition. The left arrow indicates more error than correct for fast RT, responsible for the decrease in accuracy in the left part of the CAF. The right arrow reveals another excess of errors for long RT, explaining the decrease in performance for long RT. Panel B: Example of mixture analysis for the correct distribution depicted on panel A. Although the mixture appears more clearly on errors, it is also present on correct trials as revealed by mixture analysis which indicates that the raw distribution (continuous thick line) can nicely be described as two overlapping distribution (doted lines). The characteristics of these two distributions can be estimated and analysed.

The CAF for D1 is presented on Figure 4 panel A. For compatible trials, the proportion of correct trials was rather flat, close to ceiling. In contrast, for incompatible trials, the proportion of correct responses was initially low for fast responses and monotonically increased as RT lengthens. Repeated measures ANOVA on D1 revealed a main effect of compatibility [F(1, 22) = 146.5; *p <* .0001], of quantiles [F(6,132) = 20.23; *p <* .0001] and a small interaction between the two [F(6,132) = 2.15; *p* = .052]. Trend analysis revealed that both the linear [compatible: *β*= 0.066, F(1, 132) = 15.85; *p <* .0001, incompatible: *β*= 0.141, F(1, 132) = 107.0; *p <* .0001] and the quadratic (convex) components were significant [compatible: *β*= −0.040, F(1, 132) = 5.75; *p <* .0001, incompatible: *β*= −0.041, F(1, 132) = 9.03; *p <* .0001]. Note, however, that the linear component is now positive, indicating an overall increase (although convex) of performance as RT increases. Finally on D1’s CDF (Figure 4 panel B), repeated measures ANOVA showed a trivial main effect of quantiles [F(6,132) = 256.16; *p <* .0001], of compatibility [F(1,22) = 52.84; *p <* .0001], and an interaction between the two factors [F(6,132) = 18.84; *p <* .0001]. The interaction, considered with the delta plot, indicates an increase of the compatibility effect, as RTs lengthen.

**Figure 4.**
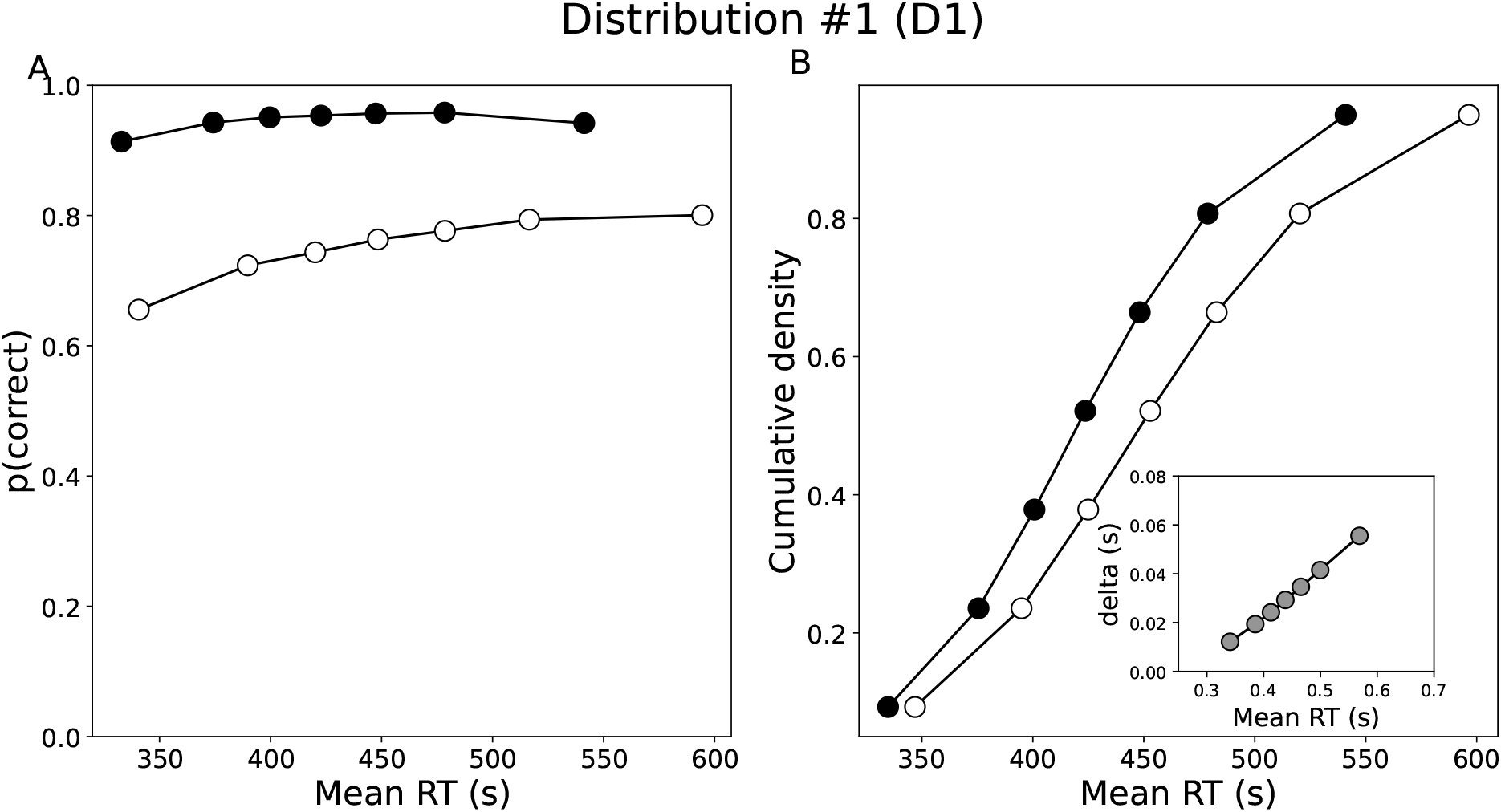
Distribution analysis of the D1 simulated distribution. A. Conditional Accuracy Function (CAF) for compatible (black circles) and incompatible (open circles) trials. Note that for incompatible trials, accuracy now increases monotonically as RTs lengthen. B. Cumulative Density Function (CDF) of the correct responses and (Inset) mean interference effect (incompatible - compatible) as function of RT. The size of the interference effect on RT increases linearly as RTs lengthen.

### DMC modeling

Attempting to fit the DMC to raw data failed, as the the fit was bad, and the model could actually not account for the data (see Figure S3). If the above reasoning is correct, only the D1 distribution should correspond to the processes that the DMC aims to model. We then fitted the model to the D1 data which should correspond to the processes that the DMC aims to model. The model very nicely fits the D1 data, as can be appreciated in Figure 5. The best fit parameters are provided on Table 1, along with the parameters obtained in previous studies in humans (adults and children). Although direct statistical comparisons are not possible, the similarities and differences in parameters deserve comments. Concerning the “controlled” route components parameters, the rate of information accumulation (*μ*) recovered in rats seems in between the one obtained in human adults and children, while the boundary for the decision to be made (*b*) seems even lower than human adults performing under speed pressure, maybe reflecting a higher impulsivity in rats. The non decision time (*Ter*) is comparable in rats and adults, and much shorter than in children. Concerning the “automatic” process parameters, the strength of the automatic activation (*A*) seems comparable in rats and in human adults and slightly lower than human children. The time course of this automatic activation (*Lat*) deserves specific comments. According to DMC, the time course of the automatic activation is critical for the between task differences in the shapes of the CDF and delta-plots. The present data indicates that rats have a much longer automatic activation than humans, be they adults or children. To further compare the time course of automatic activation across task and species, figure 6 summarizes the value of the *Lat* (normalized by the mean RT of each study) parameters obtained so far. The time course of the automatic activation is much shorter in the Simon task than in the flanker, and likely in the Stroop one (no adult data available), but does not differ betwen children and adults in the Simon task. In contrast, it is much delayed in children in the flanker (and likely in the Stroop) task compared to adults. Finally, automatic activation is much more sustained in rats, reaching values comparable to the one observed in adults in the Eriksen flanker task.

**Figure 5.**
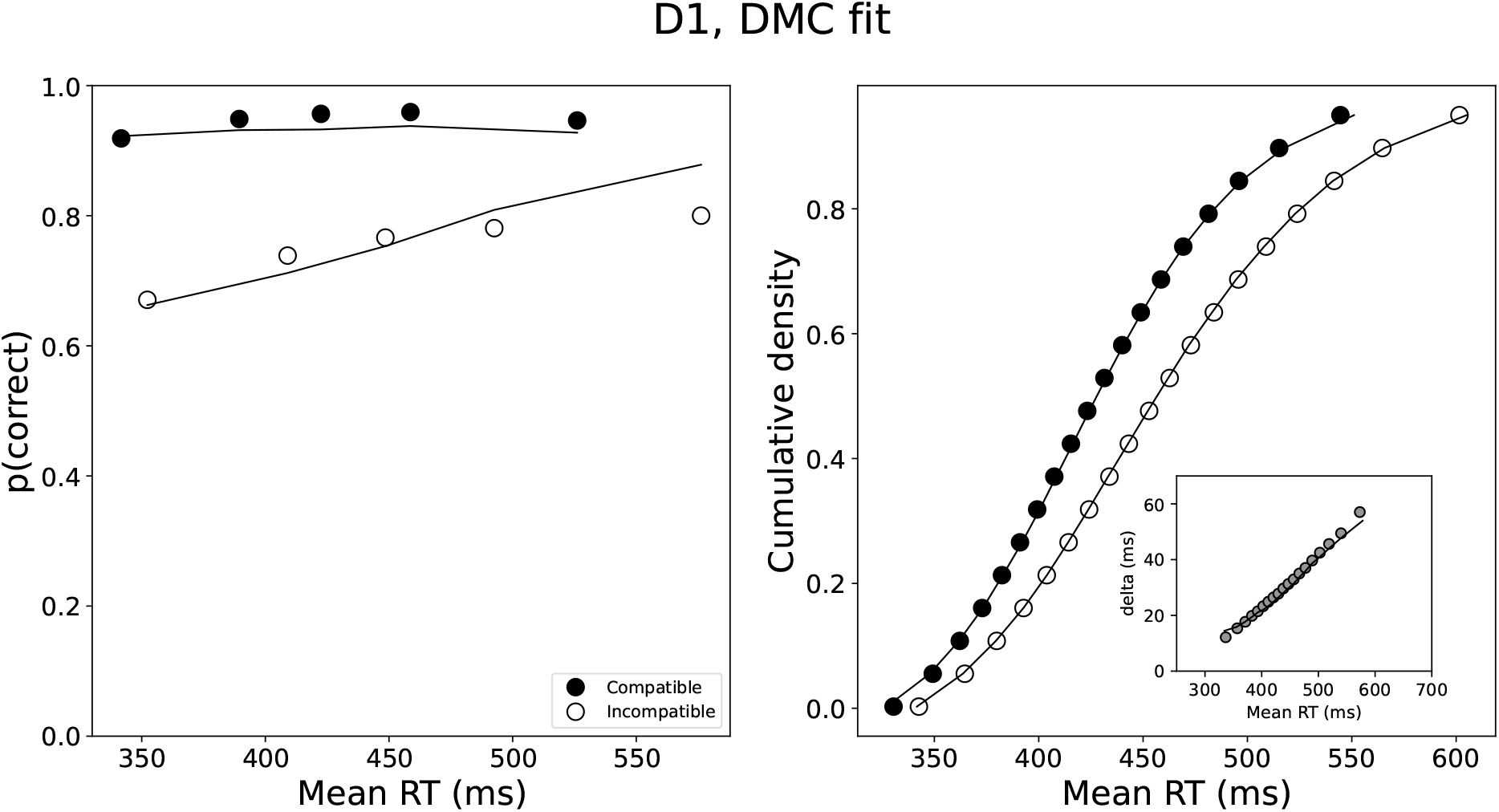
DMC fit of the D1 distribution. Circles represent D1 data (filled: compatible; open: incompatible) and lines the model fit. A. CAF B. CDF and delta plot (Inset).

**Figure 6.**
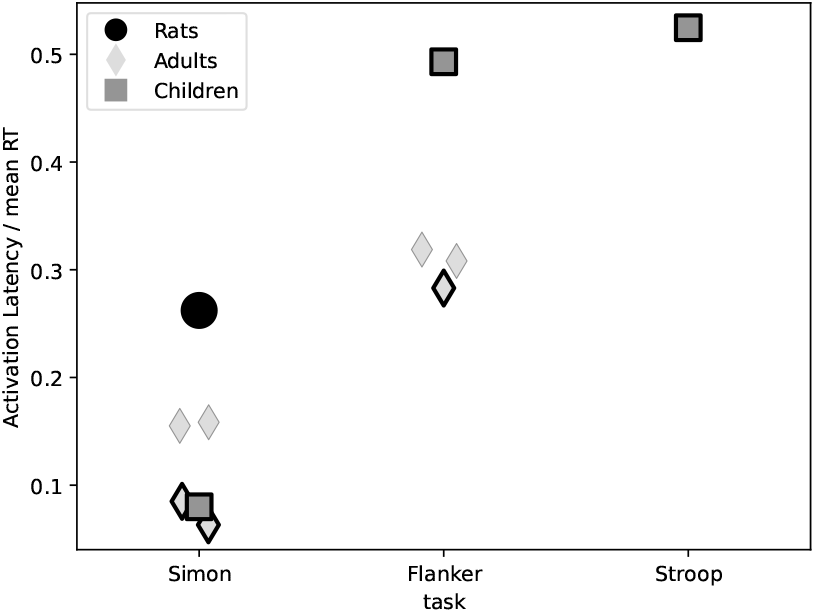
Overview of the dynamics of the automatic activation across studies in the litterature for the three main conflict tasks on which DMC has been fitted (Simon, Eriksen-flanker and Stroop). To take into account the large difference in mean RT, especially with age, the latency at which the automatic activation reaches its maximum (*A*) is divided by the mean RT. Diamonds correspond to human-adult data, squares to human-children, and the circle to the current rat data. Diamond data with a thin contour correspond to fits where the shape of the gamma function was allowed to vary. For the other ones, this value was not fitted and fixed at 2. The plotted data come from the studies listed in table 1.

We will speculate on the origin of this different time course in the discussion.

## Discussion

Being able to resist the temptation to act induced by irrelevant information is essential for adapted behavior. The Simon paradigm (along with its relatives) has been largely used to investigate these capacities in young human adults (Hommel, 2011, for an in depth analysis of the usefulness of this paradigm). Because of its simplicity, it can be used in many populations including children (Ambrosi et al., 2020; Iani et al., 2014; Smigasiewicz et al., 2020) and non-human species. Although compatibility effects have been observed on mean RT and error rates in several non-human species (Courtière et al., 2007, 2011; Czaczkes et al., 2022; Huguet et al., 2014; Nakamura et al., 2005; Urcuioli et al., 2005), the pivotal question as to whether they stem from the same underlying processes remains open: if the compatibility effect is driven by the same processes along the phylogeny (down to a certain point/species), any explanation of the effect, including in humans, would have to be compatible with the currently known cognitive development of the other species. A core characteristic of the compatibility effect in humans is its dynamics (De Jong et al., 1994; Pratte et al., 2010; Ridderinkhof, 2002) which has never been studied in rodents. Observing similar dynamics in both species would strengthen the idea that same processes are at stake, while very different dynamics would speak against this view. To advance on this comparison, we analysed the shape of correct RT distribution on CDF, the dynamic of accuracy as a function of RT on CAF and we fitted a recent model of conflict tasks, in a newly developed version of the Simon task in rats. Contrary to what is observed in human (visual versions), CDF analysis reveals that the difference in RT between compatible and incompatible trials increase as RTs lengthen. Recent theoretical work (Ulrich et al., 2015) directly relates this pattern to the dynamics of incorrect response activation (potentially including its suppression). The monotonic increase in the compatibility effect as RT lengthens suggests a long lasting activation of the automatic response. This view is supported by the CAF analysis which revealed clear similarities but also noticeable differences with performance of human adults. First, a decrease in accuracy was observed for the fastest RT on incompatible trials comparable to the pattern observed in humans, likely reflecting an automatic activation of the response ipsilateral to the stimulus position. But a marked difference appeared for the long RTs: while accuracy converges to almost perfect as RT lengthens in humans, it dramatically decreases in rats. This decrease was apparent in both compatible and incompatible trials and explains the globally slower RT for errors observed on mean performance. While unexpected, this pattern is not unique. Although they did not compute CAF, a close examination of Kaneko, Tamura, Kawashima, and Suzuki (2006)’s data suggests similar mechanisms in a different type of conflict task in rats: in a Stimulus-Response compatibility task (Fitts & Deininger, 1954), “short-RT” errors (which would induce a drop of accuracy for fast responses) were followed by “long-RT” errors (see their figure 4), that would induce a decrease in the CAF for the slowest RT. Interestingly, a similar pattern can also be observed in young children. In a (complex) Stroop task, Bub, Masson, and Lalonde (2006) report that 7 to 11 years-old children present a drop of accuracy for fast responses, but also a large decrease for slow RTs, for both compatible and incompatible trials (see their figure 3a), as observed in the present report. Bub et al. speculated that the non-specific decrease in accuracy for long RT might reflect a deficiency in maintaining the task goal. In the present report, mixture analysis helped in clarifying this pattern. It revealed that measured RT actually stem from two different distributions with different characteristics. While a first distribution (discussed below) has all the classical characteristics of regular RT distribution (fast, asymmetric etc.), the second one is much slower and has a more normal shape (more detailed analysis are provided on Supplementary material). This second distribution is at the origin of the sharp decrease in accuracy as RT lengthens in the raw data. Although speculative, it could be that a similar secondary distribution was present in Bub et al. (2006) data explaining their pattern of results in children. It could reflect a loss of the task goal as rats take time to respond, as has been proposed in children (Bub et al., 2006; Diamond & Taylor, 1996). This view is supported by increasing proportion of trials belonging to D2 at the end of the sessions (see Supplementary Material). If such is the case, adding a task-reminder (see Barker & Munakata, 2015, for an example in children with a Go/No-go task) should decrease the proportion of the second distribution.

The first distribution extracted by the mixture analysis likely captures the “true” RT distribution and should hence be more comparable to regular human data. Supporting this view, this distribution was nicely fitted by the DMC, and CAF constructed based on this first distribution were much closer to the one observed in human adults: on incompatible situation, accuracy monotonically increases as RT lengthens, although 100% accuracy was never reached. Note also that the increase in accuracy is rather “slow” (small and regular increase from one quantile to the next). This slow increase in accuracy is captured by a delayed peak of the automatic activation (around 130 ms, compared to around 30-60 ms even in children) estimated from the DMC fit. This is reminiscent of the results obtained in children performing a Stroop task (Ambrosi et al., 2019, 2020): while CAF of 5-years old children did not converge to 100%, this gap decreased with age, until reaching almost perfect accuracy at 12 years old onward. Overall, the data suggest a long lasting automatic activation in rats performing a “simple” Simon task, similar to the one observed in children performing more demanding conflict tasks. The question as to whether this long-lasting activation reflects a slow suppression mechanism in rats and young children (compared to human adults), is a tentative hypothesis that needs to be explicitly tested, opening new perspectives.

In conclusion, advanced data processing techniques, coupled with distribution analyses, mixture analysis and model fit in conflict tasks, allowed to reveal similarities and differences between humans (adults and children) and rodents, establishing bridges between phylogenetic and ontogenetic cognitive development. It appears that rats tend to respond according to two strategies, one very close to the one observed in humans (especially close to children performing a more complex task), while the nature of the second (which might also be present in young children) still needs to be better specified. These results open the possibility to quantitatively evaluate and compare the onto- and phylo- genetic evolution of cognitive control.

## Supporting information

Supplementary results

## Footnotes

1 The same fits were performed with the DMCfun R package by the same authors. The results were the same.

