## Supplementary results for "At the roots of interference control: Conflict task in rats reveals the commonalities of onto- and phylo-genetic development"

Supplementary material

Julien Poitreau, Frédéric Ambroggi, Thierry Hasbroucq, Francesca Sargolini and Boris Burle

LNC, Aix-Marseille Université, CNRS

At the roots of interference control:

Conflict task in rats reveals the commonalities of onto- and phylo-genetic development

Supplementary material

### Supplementary Methods

#### Apparatus

Behavioural procedure took place in 8 operant chambers (Med-Associates) of 29×24×29 cm (inner perimeter) with a grid as the floor, 2 side walls and ceiling in transparent acrylic glass, and front and back walls in stainless steel. The feeder (5×5×2 cm) was connected to a pellet dispenser, attached behind the back wall of the cage. The front wall was curved and equipped with 5 nose-pokes (NP), consisting in squares of 2.5 cm, 1 cm deep and positioned 2 cm above the floor. Only 3 NP were used, the central one and the 2 directly lateral on each side. The 2 others were blocked with a metal cap. All the NP were equipped with a light bulb inside to provide visual stimuli and infrared photocells allowing to detect animals responses. Except for the light of the visual stimuli, the chambers were completely in the dark during the whole sessions. A transparent acrylic wall with a 10×8 cm door was positioned 12 cm from the front wall to constraint animals position in front of the central NP. A loud-speaker attached to the ceiling of the cage provided auditory feedback (low-pitch or high pitch). Each chamber was in a sound-attenuating box, ventilated by a low-level noise fan. Cages were controlled by a computer and Med-Associates system, with programs wrote in Med-State Notation. An infrared camera attached to the ceiling of the cage was used to monitor and record animals behaviour.

All events during the task (i.e. stimuli, pokes, rewards) were recorded with Med-Associates system and were extracted with MedPC-to-Excel utility and processed with Python and R scripts (available at xxx).

#### Task shaping

Animals were trained 30 minutes, 2 times a day, 5 days a week, with approximately 4h between sessions. Training was composed of 3 consecutive phases.

1. Getting used to the operant chamber

   - Feeder habituation: a sucrose pellet was given at each detection of the rat in the feeder (2 days).
   - Central NP-feeder association: Central NP light switch on indicated trial start. A poke in the central NP then triggered a sucrose pellet in the feeder and light switch off. Inter-trial interval

was set to 5 s. At least 30 engaged trials and systematic retrieval of the reward were required to move on to the next learning step

2. Learning the response rule:

   - Learning to use the lateral NPs: After the central poke, the response to give was indicated by a **bilateral** stimulus in the lateral NPs. The response rule (right or left NP) was given by the light intensity (bright or dim). Rule was counterbalanced. A correct response triggered a sucrose pellet. There was no time limit to give a response after the initial central poke. During this first rule learning step, only one type of stimulus (bright or dim) was presented during the session, and sessions were performed in alternation. At least 2 sessions with 30 engaged trials and systematic lateral poke was needed to go to the next learning step.
   - Mixing responses: Same procedure as before with both light stimuli (dim and bright) presented in the same session in a randomized manner. After an error, a 5 s punishment delay was added and the same type of trial was restarted to avoid stereotyped responses. The learning criterion was fixed at 85% of correct rate.

3. Final adjustments of the task.

   - Speeding up responses: In humans, reaction time is defined as the time needed to give a response as fast and as accurate as possible. With the aim to get closer to this definition, rats behaviour were speeded-up by reducing, first the response window to 1.5 s after the central poke and then the stimuli presentation to 500 ms. Omissions were considered errors and treated the same way.
   - Preparatory period: In order to stereotype response procedure with the aim to have a more accurate measure of reaction time, we wanted to ensure that the animals stay in a central position when the side stimuli were presented. For this purpose, we added to the chamber an acrylic door, 12cm in front of the NPs to restrain animals position and trained them to hold the central NP for 1 sec before the stimulus presentation. To do so, stimulation delay after the central poke was progressively incremented by 50 ms every 5 consecutive successful trials until reaching 1 sec.
   - Unilateral stimulus: The spatial interference was introduced by modifying the stimulus, which no longer appeared bilaterally (i.e. in both lateral NPs) but unilaterally in a pseudo-randomized manner. As we were not interested in the learning dynamic, rats were trained with unilateral stimuli until mean performances reach a plateau. After what, the test phase began and behaviour was recorded during 16 sessions for analysis.

In case of several good responses in a row, stimulus position and intensity were pseudo-randomly distributed to prevent more than three repetitions of the same type of trial. Sessions were 30min long or ended after animals collected a maximum of 150 pellets. As we were not interested in the learning dynamic, rats were trained until performance plateau before the test phase, consisting in 16 sessions (8 consecutive days) during which performances were recorded for analysis.

### Supplementary Results

#### Mixture analysis

**Comparing single distribution and mixture.** For each rat, compatibility and accuracy, we compared two models: one with only one distribution (“single”) and one as a mixture of two distributions (“mixt”). Out the 92 elementary cells, the mixture model has a lowest AIC for 80 cells, and only 12 of them favored the single model. To better asses the evidences for and against the mixture, we compared the difference in AIC by taking the absolute value of the difference. Since the difference in AIC is small when the single distribution is favored, we choose to use the mixture model, even when the single distribution was (slightly) favored.

**Properties of D2.** Overall, D2 was much longer than D1 (compare the x-axis of main Fig. 4 and of Fig. S2).

***CAF.*** An ANOVA performed on CAF with quantiles and compatibility as factors revealed a main effect of quantiles ($F(6,132) = 18.46$; $p < .001$), a main effect of compatibility ($F(1,22) = 24.31$; $p < .001$), but not interaction between the two ($F(6,132) = 0.18$). Trends analysis revealed that for the two compatibility conditions, both the linear (decreasing) [Compatible: $\beta$= -0.156, $F(1, 132) = 34.17$; $p < .0001$, incompatible: $\beta$= -0.178, F(1, 132) = 7O.84; $p < .0001$] and the quadratic (convex) components were significant [Compatible: $\beta$= - 0.087, $F(1, 132) = 10.48$; $p < .0001$, incompatible: $\beta$= -0.058, $F(1, 132) = 7.57$; $p < .01$].

***CDF.*** On CDF, a main effect of compatibility showed up ($F(1,22) = 17.02$; $p < .001$), along with a trivial main effect of quantiles ($F(6,132) = 192.58$ ; $p < .001$) and, more importantly, an interaction between these two factors ($F(6,132) = 16.15$ ; $p < .001$). The interaction, in accordance with the delta-plots (see Fig. S2, panel B, inset), confirms the interference effect increases as RT increases.

***Evolution of D2 proportion within sessions.*** Following Bub et al., we hypothesized that D2 could be attention related and stem for a lost of task goal. If so the proportion of D2 should increase as rats become more fatigued/less motivated, that is towards the end of the sessions. We hence examined the evolution of D2 proportion between the first 5 and last 5 minutes of the sessions for each trial type (compatibility and correctness). Proportion of D2 in these subsets of data were calculated with mixture

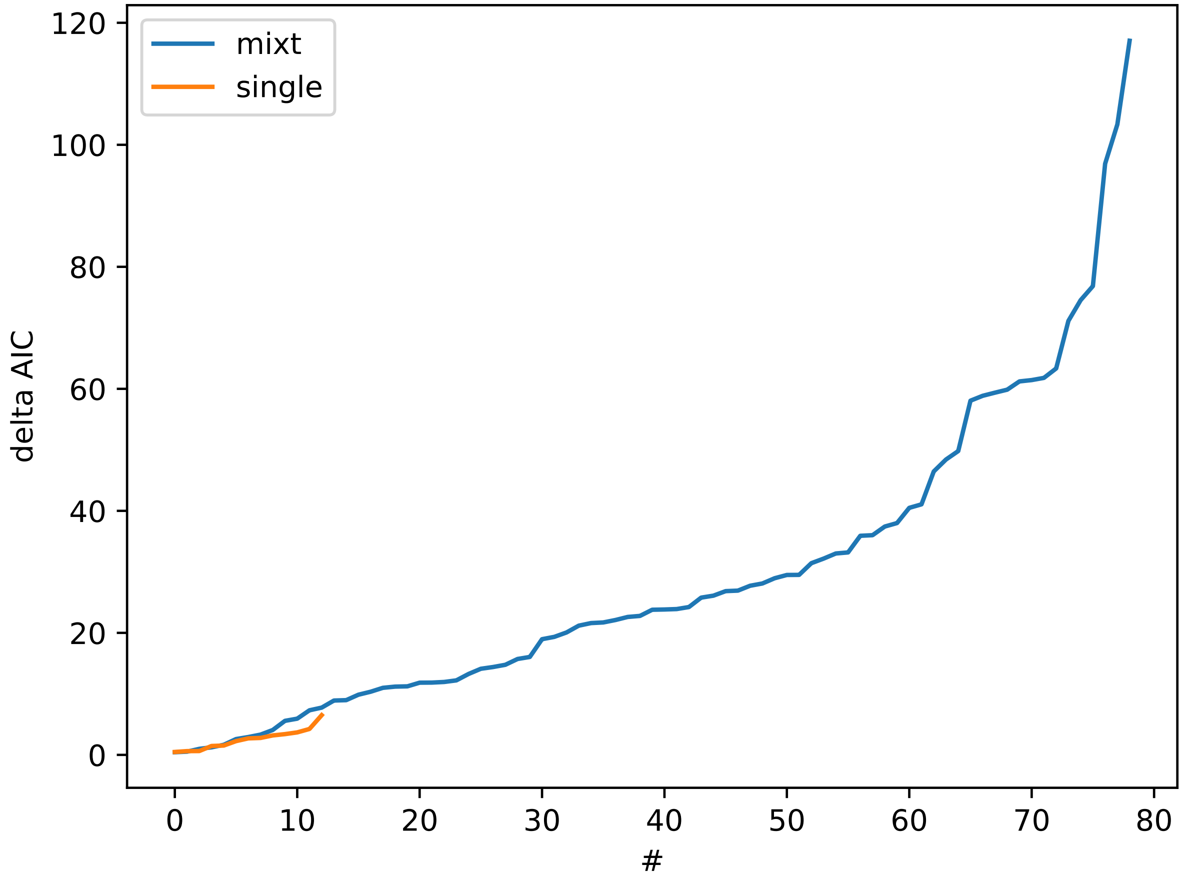


*Figure S1*. Sorted difference in AIC for the case where the mixture was favored (blue) and when the single distribution was (orange). While the difference in AIC is pretty large when the mixture is favored, the differences are very small when the single is.

analyses, the same way as previously described. Two rats in the first 5 minutes and 4 in the last 5 minutes of the 16 sessions did not perform any compatible erroneous trial. To keep these rats into the analyses despites those few missing data, we input them with the median proportion of this trial type for each 5 minutes bin to neutralize them. Mean D2 proportions for each trial type in each 5 minutes bin is displayed in Table 1. A repeated measures ANOVA was performed on these proportions with compatibility, correctness and time bins as within-subject factors. This analysis revealed main effects of compatibility [F(1,22) = 22.88; p < .0001] and correctness [F(1,22) = 129.96; p < .0001]. These two factors interact [F(1,22) = 24.48; p < .0001] and so do time bins and correctness [F(1,22) = 7,62; p < .01]. As we were interested in the difference in D2 proportion between time bins we performed a one-sided pairwise T-test to explore this last interaction, which revealed a significant increase of the D2 proportion between the first 5 and last 5 minutes of the session for incompatible erroneous trials [t = -3.72; df = 22; p < .001], but not on

Table 1

*Evolution of the D2 proportion during the sessions.*

| | Compatibles | | | |
|---|---|---|---|---|
| | Corrects | | Errors | |
| | M | CI | M | CI |
| D2 proportion % | | | | |
| First 5 mn | 22.05 | [21.08,23.02] | 54.15 | [52.15,56.14] |
| last 5 mn | 17.59 | [16.59,18.59] | 59.02 | [57.13,60.90] |

| | Incompatibles | | | |
|---|---|---|---|---|
| | Corrects | | Errors | |
| | M | CI | M | CI |
| D2 proportion % | | | | |
| First 5 mn | 21.68 | [20.49,22.87] | 26.46 | [24.97,27.94] |
| last 5 mn | 19.32 | [18.62,20.03] | 38.92 | [37.73,40.11] |

compatible errors. Although the larger proportion of D2 on compatible may seem counter-intuitive, it likely simply reflect the fact that the **number** of D2 trials is comparable on compatible and incompatible trials. However, the number of “regular” error being lower on compatible trials, the **proportion** of D2 errors is higher.

### DMC modeling of raw data

The DMC model (Ulrich, Schröter, Leuthold, & Birngruber, 2015) was fitted to the the raw data (see Fig. S3). As could be expected, the model cannot account for the decrease in accuracy as RTs lengthen, leading to a large misfit for the long RT, for both compatible and incompatible trials in the CAF (Fig. S3A). The fit on the RT distribution is also pretty bad, with large deviations on the delta-plots (Fig. S3B, inset)

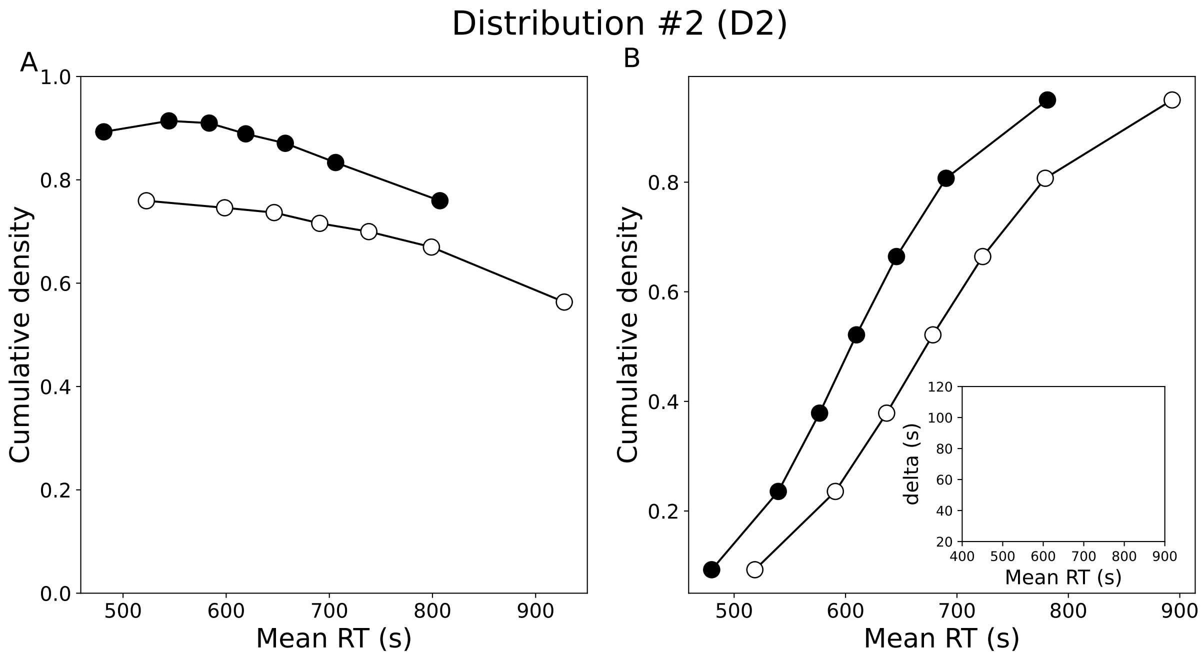


*Figure S2*. CAF and CDF for D2

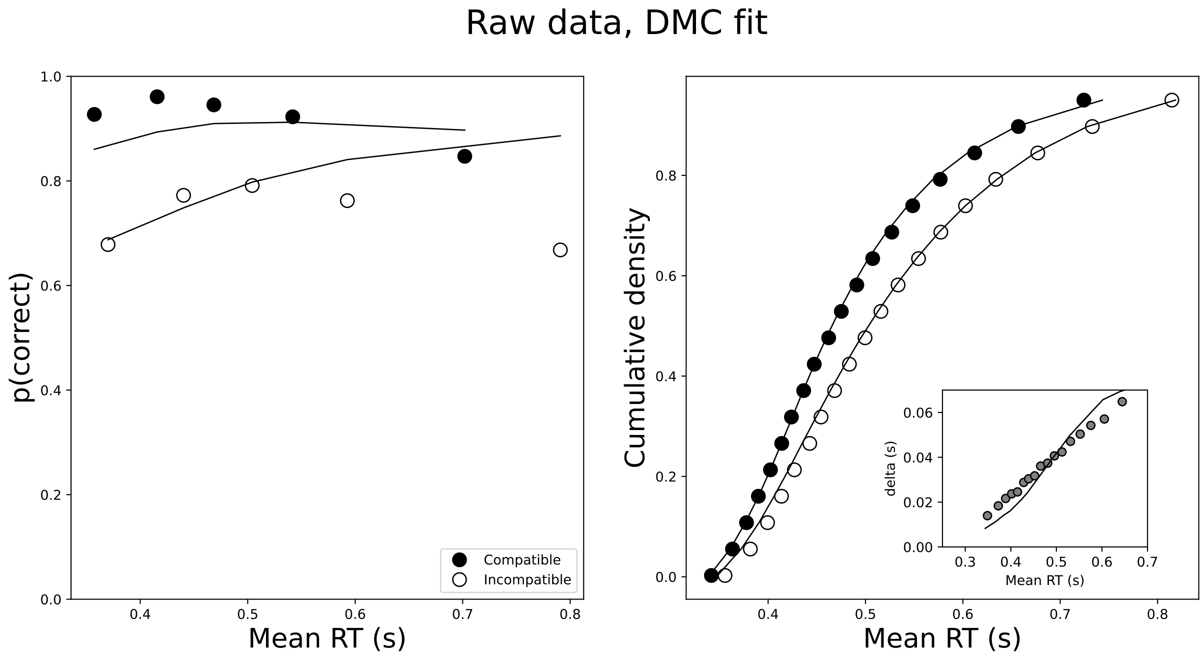


*Figure S3*. Fit on raw data
